# Compartmentalization of enhanced biomolecular interactions for high-throughput drug screening in test tubes

**DOI:** 10.1101/2020.01.19.911149

**Authors:** Min Zhou, Weiping Li, Jian Li, Leiming Xie, Rongbo Wu, Liang Wang, Shuai Fu, Wei Su, Jianyang Hu, Jing Wang, Pilong Li

## Abstract

Modification-dependent and -independent biomolecular interactions (BIs), including protein-protein, protein-DNA/RNA and protein-lipid, play crucial roles in all cellular processes. Dysregulation of BIs or malfunction of the associated enzymes results in various diseases, thus they are attractive targets for therapies. High-throughput screening (HTS) can greatly facilitate the discovery of drugs for these targets. Here we describe a HTS drug discovery method, called <u>c</u>ompartmentalization of <u>e</u>nhanced <u>b</u>iomolecular interactions in test tubes (CEBIT). CEBIT uses selective recruitment of biomolecules into phase separated compartments harboring their cognate binding partners as readouts. CEBIT were tailored to detect various BIs and associated modifying enzymes. Using CEBIT-based HTS assays, we successfully identified known inhibitors of the p53/MDM2 interaction and of SUV39H1 from a compound library. CEBIT is simple and versatile, and is likely to become a powerful tool for drug discovery and basic biomedical research.

## Introduction

Complex networks of modification-dependent and -independent biomolecular interactions (BIs) lay the foundation of diverse cellular processes including DNA replication, transcription, protein translation, and signal transduction. Dysregulation of BIs causes many diseases, such as neurodegenerative diseases, infection and cancers^1, 2^. Hence, aberrant BIs and the associated modifying enzymes, if any, are attractive targets for therapeutic drug discovery^3–5^.

High-throughput screening (HTS) provides a powerful strategy for the discovery of drugs that modulate various BIs^6^. Many HTS technologies are widely used, such as surface plasmon resonance (SPR), fluorescence polarization (FP) and fluorescence resonance energy transfer (FRET). These methods all have major limitations. Many factors can interfere with the outcomes of SPR, such as ionic strength, DMSO content, charges of immobilized proteins, and nonspecific binding to the biosensor^7–10^. FP requires not only a big difference in molecular mass between the fluorescent probe and the target protein, but also high affinity probe to achieve large signal window^9, 11^. These drawbacks largely limit its application. FRET, a third commonly used HTS method, has a narrow signal window and high background due to the limited energy transfer distance (<10 nm)^9, 11, 12^. It is vital to develop more robust and easily implementable HTS technologies for efficient drug discovery.

Cells use numerous membrane-enclosed and membraneless organelles to compartmentalize biochemical reactions. Membraneless organelles, such as P granules^13^, nucleoli^14^, stress granules^15^, and post synaptic densities^16^, are collectively referred to as biomolecular condensates. These condensates are assembled via liquid-liquid phase separation (LLPS) driven by multivalent interactions^17, 18^. Generally, the constituents of various condensates are divided into two groups: one group is composed of scaffolds, which are essential for the assembly of these condensates and the other group is composed of clients, which are dispensable for the formation of the condensates but can partition into these condensates via directly or indirectly binding to the scaffolds^19^. Inspired by the scaffold-client model, we explored the possibility of using a similar architecture to implement a system for assaying modification-dependent and -independent BIs, which can be used for efficient discovery of drugs targeting these BIs and associated enzymes.

Our system consists of two parts: (1) the scaffold, which drives the formation of phase separated compartments; and (2) the BI of interest, of which one partner is fused with the scaffold to generate a composite scaffold and hence automatically enriched in the compartments and the other partner (defined as client) is recruited to the compartments via specific interaction. For visualization, the composite scaffold and the client are labeled with different fluorescent probes such as GFP and mCherry. Thus, the BI of interest is assessed by the enrichment of mCherry fused client within the GFP labeled phase separated compartments. This design can also be used to measure the activities of enzymes modifying biomolecules.

This strategy is called <u>c</u>ompartmentalization of <u>e</u>nhanced <u>b</u>iomolecular interactions in test tubes (CEBIT). We also developed and evaluated a similar method for studying protein-protein interactions (PPIs) in cells (see the accompanying manuscript). We extensively validated CEBIT using some representative PPI pairs. Subsequently, we tailored CEBIT to probe molecular modifications and the associated enzymes.CEBIT is very amenable for HTS. Using the MDM2/p53 interaction and the SUV39H1-mediated H3K9 methylation reaction as two test cases, we successfully identified inhibitors of the MDM2/p53 interaction and of SUV39H1 from a commercial compound library. CEBIT is a simple and versatile method, which will have many applications in basic and translational research.

## Results

### Establishing robust multivalent scaffolds

First, we sought to establish some multivalent scaffolds that can robustly drive the formation of compartments via LLPS. Numerous studies have demonstrated that interactions between linear and/or dendrimeric (branched) multivalent modular proteins can result in LLPS^17, 19–22^. The *Saccharomyces cerevisiae* protein SmF (referred to as SmF hereafter) is known to form a stable tetradecameric (referred to as 14-meric for simplicity hereafter) complex upon expression alone in bacteria^23^. We tested whether it was possible to reliably achieve dendrimeric multivalence of various domains/motifs when they were fused to SmF. We created two fusion proteins, one with green fluorescent protein (GFP) fused to the C-terminus of SmF (SmF-GFP) and the other with the second Src homology 3 (SH3) domain of human NCK1 fused to the C-terminus of SmF-GFP (SmF-GFP-SH3). Size-exclusion chromatography coupled with multi-angle light scattering (SEC-MALS) analysis indicated that SmF-GFP also formed a 14-meric complex in solution, and further fusion of a SH3 domain to SmF-GFP did not alter the 14-meric status (Fig. 1a). Therefore, we concluded that SmF is a robust 14-meric protein for multimerizing protein domains/motifs.

**Figure 1.**
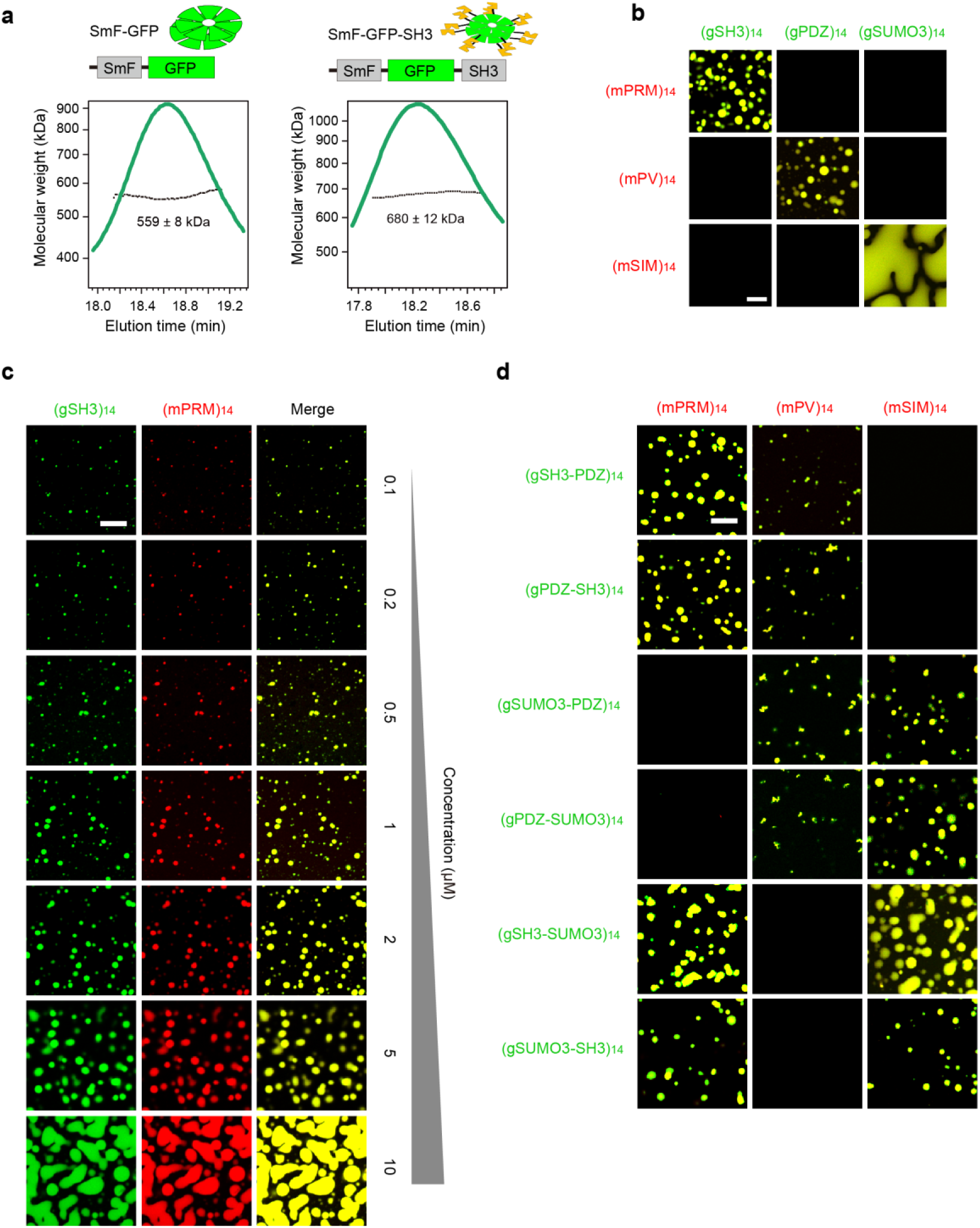
Phase separation of multivalent scaffold proteins. (a) Multimerization of module domains by fusion with a tetradecameric (referred to as 14-meric for simplicity hereafter) protein, yeast SmF. GFP is fused to the C-terminus of SmF and the resulting protein SmF-GFP (theoretical molecular mass 566 kDa) is 14-meric based on SEC-MALS experiments. SH3 is then fused to the C-terminus of SmF-GFP and the resulting protein SmF-GFP-SH3 (theoretical molecular mass 671 kDa) is also 14-meric based on SEC-MALS experiments. (b) Phase separation of multimerized protein protein interactions. Domain structures of the model protein pairs used here are shown in Supplementary Figure S1a. (gSH3)_14_, (gPDZ)_14_, and (gSUMO3)_14_ were cross-mixed with (mPRM)_14_, (mPV)_14_, and (mSIM)_14_ for assessment of phase separation. Merged images are shown. All proteins are at 5 µM. (c) Phase separation assays of (gSH3)_14_ and (mPRM)_14_ over a range of protein concentrations. Individual and merged images are shown. (d) Binary combinations of SH3, PDZ, and SUMO3 were fused with SmF-GFP to generate 6 composite scaffold proteins (Supplementary Figure S1b). These six proteins were mixed with (mPRM)_14_, (mPV)_14_, or (mSIM)_14_ at 1 μM. Merged images are shown. All scale bars, 10 μm.

Similarly, we investigated the *Bacillus subtilis* protein Hfq (BsHfq), which is known to form a stable hexameric complex^24^. We confirmed that BsHfq can reliably achieve dendrimeric multivalence of fused domains/motifs (Fig. S1).

### Multimerized PPIs undergo LLPS

Next, we investigated whether multimerized PPIs, created by fusion to SmF, could mediate LLPS. We selected three model PPI pairs: (1) the second SH3 domain of human NCK1 and the proline-rich motif (PRM) of DLGAP2^17^; (2) the third PDZ domain of human PSD95 and its ligand, KKETPV (abbreviated to PV)^25^; and (3) SUMO3 and the SUMO3 interacting motif (SIM)^19^. In each PPI pair, one partner was fused to SmF-GFP and the other was fused to SmF-mCherry (Fig. S2a). The resulting proteins are abbreviated as (g/mDOMAIN/MOTIF NAME)_14_ for simplicity, in which g/m stands for GFP/mCherry and the subscript 14 represents the 14-meric status of the protein complexes. We carried out cross-mixing reactions between (gSH3)_14_, (gPDZ)_14_, or (gSUMO3)_14_ and (mPRM)_14_, (mPV)_14_, or (mSIM)_14_. All three cognate multivalent PPI pairs, but not the non-interacting PPI pairs, gave rise to marked LLPS (Fig. 1b). We next carried out phase-separation assays of (gSH3)_14_ and (mPRM)_14_ at 1:1 molar ratio over a large module concentration range, 100 nM to 10 μM. Numerous droplets formed at module concentrations as low as 100 nM (Fig. 1c). We further fused all binary combinations of the three modules, SH3, PDZ, and SUMO3, to the C-terminus of SmF-GFP to yield six composite scaffolds, in which two different interacting domains are present in two different orders (Fig. S2b).

Cross-mixing the six proteins with (mPRM)_14_, (mPV)_14_, or (mSIM)_14_ showed that LLPS occurs as long as and only if a cognate multivalent PPI pair is present (Fig. 1d). These results suggest that the presence of a non-cognate binding module does not appreciably affect the ability of multivalent PPI to drive LLPS. Collectively, these data further revealed the robustness of SmF as a multivalent scaffold to multimerize PPIs for inducing LLPS.

### Recruitment of clients into phase-separated compartments via specific PPIs

As revealed by Banani et al., clients are recruited into scaffold-induced condensates by binding to free sites of scaffolds^19^. We wondered whether PPIs of interest could be studied similarly as clients. To test this, we used (gSH3-PDZ)_14_ (abbreviated for SmF-GFP-SH3-PDZ, Fig. S2c) and (gPRM)_14_ to drive LLPS, presumably via the multivalent SH3/PRM interaction, and we simultaneously used mCherry-fused PDZ ligand, KKETPV (abbreviated to mPV, Fig. S2c), as a client to assess the interaction between PDZ and KKETPV (Fig. 2a). (gSH3-PDZ)_14_ and (gPRM)_14_ formed green compartments (Fig. 2b, lower panels). mPV was recruited into these droplets, as shown by the enriched mCherry signal (Fig. 2b, KKETAV = 0 μM). To test whether the recruitment is reversible, we used a peptide, KKETAV, which has a slightly higher affinity for PDZ than KKETPV^25^, to compete with mPV. The enrichment of mPV was gradually reduced by increasing concentrations of the competing peptide (Fig. 2b, red), which was confirmed by quantification of mCherry signal in the green droplets (Fig. 2c). These results indicated that mPV was recruited into droplets via binding to PDZ.

**Figure 2.**
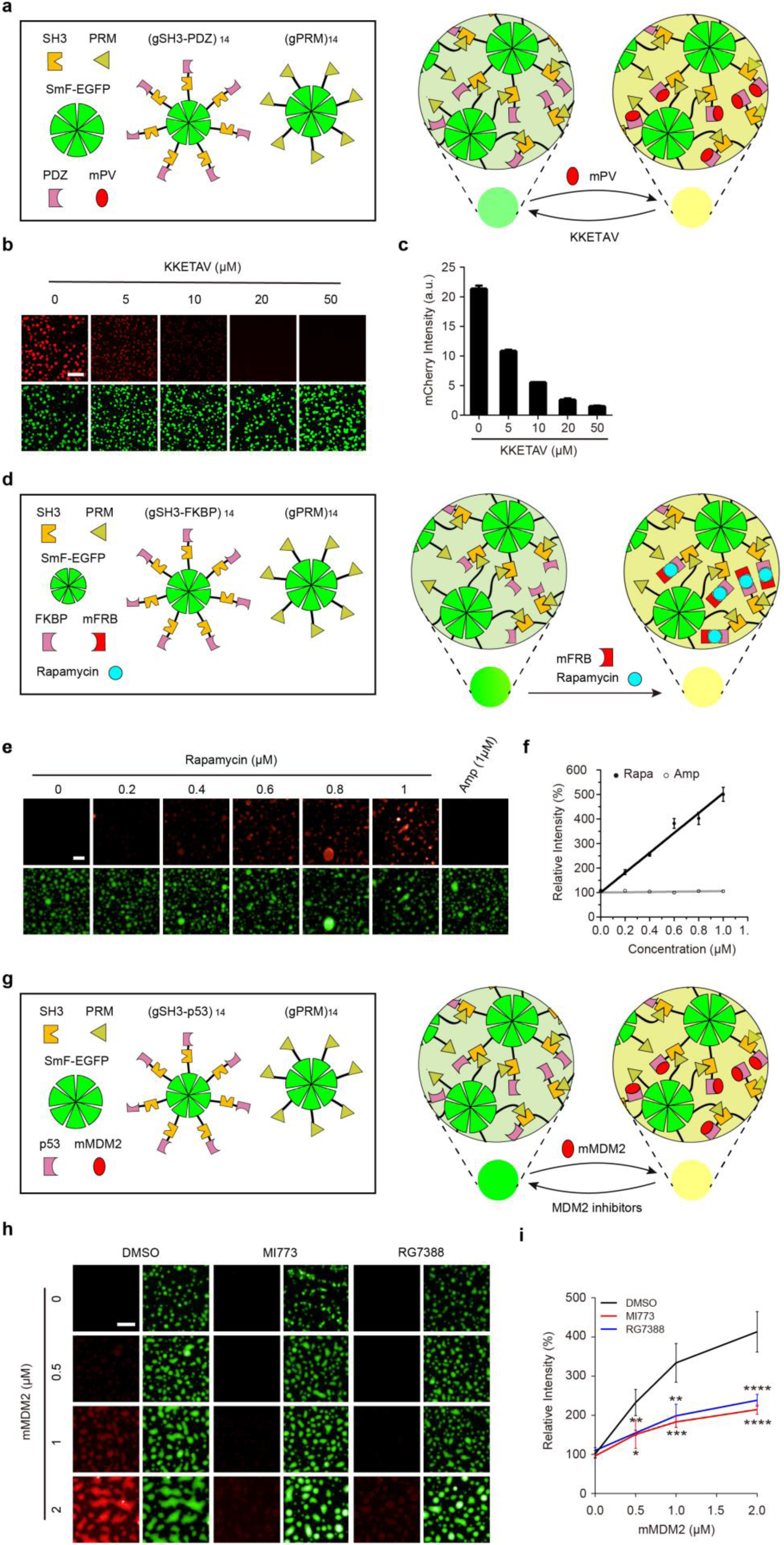
Recruitment of client proteins into phase-separated compartments via specific interaction. (a) Schematic diagram of the CEBIT based assay to assess the interaction between the PDZ domain and its ligand KKETPV. The individual modules are shown in the left box, together with the two multimeric scaffold proteins, (gSH3-PDZ)_14_ and (gPRM)_14_, that induce LLPS. The mCherry-fused PDZ ligand, KKETPV (mPV), is shown in red. The right panel shows the phase-separated compartments formed by the scaffold proteins. mPV partitions into the compartments by interacting with the PDZ module. Binding of mPV to PDZ is prevented by a competitive inhibitor, KKETAV. (b-c) mPV (5 μM) was recruited into phase-separated compartments induced by (gSH3-PDZ)_14_ and (gPRM)_14_ (1 μM each). The recruitment was inhibited by KKETAV. Representative fluorescence images (b) and quantification of mCherry signal in the droplets (c) are shown. (d) Schematic diagram of CEBIT to study the interaction of FKBP-rapamycin-FRB ternary complex. The two multimeric scaffold proteins, (gSH3-FKBP)_14_ and (gPRM)_14_, induce phase separated compartment. The mCherry-fused FRB (mFRB) as the client, partitions into the compartments via the mediation of rapamycin. (e-f) Rapamycin-induced recruitment of mCherry-fused FRB (mFRB, 1 μM) into phase-separated compartments formed by (gSH3-FKBP)_14_ and (gPRM)_14_ (1 μM each). Ampicillin was used as the control. Representative fluorescence images (e) and quantification of mCherry signal in the droplets (f) are shown. All mCherry signals were normalized to that without addition of small molecules. (g) Schematic diagram of CEBIT to study the p53/MDM2 interaction. The two multimeric scaffold proteins, (gSH3-p53)_14_ and (gPRM)_14_, induce phase separated compartment. The mCherry-fused MDM2 (mMDM2) as the client, partitions into the p53 enriched compartments but is reversed by MDM2 inhibitors. (h-i) Various concentrations of mMDM2 were recruited into phase-separated green droplets formed by (gSH3-p53)_14_ and (gPRM)_14_ (1 μM each). The recruitment was suppressed by the MDM2 inhibitors MI773 and RG7388 (5 μM each). Representative fluorescence images (h) and quantification of mCherry signal in the droplets (i) are shown. mCherry signal was normalized to that without addition of mMDM2. * 0.0332< p < 0.1234; ** 0.0021< p < 0.0332; *** 0.0001< p < 0.0021; and **** p < 0.0001. All scale bars, 10 μm.

FKBP/FRB is a well-characterized high-affinity interaction that is mediated by rapamycin^26^. We wondered whether our system can assay similar interactions. We generated a composite scaffold (gSH3-FKBP)_14_ (abbreviated for SmF-GFP-SH3-FKBP) and the corresponding client mFRB (abbreviated for mCherry-FRB) (Fig. 2d and S2c). (gSH3-FKBP)_14_ readily underwent phase separation with (gPRM)_14_. Additionally, in the presence of increasing concentrations of rapamycin, but not the negative control ampicillin, recruitment of mFRB into the green droplets increased in parallel, as indicated by the enhanced mCherry signal (Fig. 2e-f).

Subsequently we focused on two well-known cancer-related targets, MDM2 and XIAP. MDM2, an endogenous inhibitor of the tumor suppressor protein p53, binds to the N-terminal transactivation domain of p53 and mediates its degradation^27, 28^. XIAP is a suppressor of the proapoptotic protein caspase-9. The suppression is alleviated by the binding of Smac to the Bir3 domain of XIAP^29^. Importantly, potent inhibitors of MDM2 and XIAP have been developed^5, 30, 31^. We chose MDM2/p53 and Smac/XIAP to assess the feasibility of using CEBIT for drug discovery. The MDM2-binding region of p53 was fused with (gSH3)_14_ to generate the composite scaffold (gSH3-p53)_14_ and MDM2 was labeled with mCherry (abbreviated to mMDM2) as the client (Fig. 2g and S2c). As the concentration of mMDM2 increased, its enrichment in the green droplets formed by (gSH3-p53)_14_ and (gPRM)_14_ was significantly increased (Fig. 2h-i). This enrichment was significantly reduced by two potent MDM2 inhibitors, MI773 and RG7388 (Fig. 2h-i). Similarly, the XIAP-binding region of Smac was fused to the N-terminus of SmF-SH3 to generate (Smac-SH3)_14_ (abbreviated for Smac-SmF-SH3) as a composite scaffold and the Bir3 domain of XIAP was tagged by SNAP as the client (Fig. S2d). Phase separation of (Smac-SH3)_14_ and (gPRM)_14_ formed green droplets, into which SNAP-Surface 546-labeled SNAP-XIAP was recruited (Fig. S3). This recruitment was blocked by the XIAP inhibitor GDC-0152 (Fig. S3). These data demonstrate that CEBIT is competent for studying various PPIs by conjugating one member of the target PPI to a phase-separated scaffold and the other to a fluorescent label as the client.

### A high-throughput screening assay targeting the MDM2/p53 interaction

Next, we asked whether CEBIT can be used for drug discovery via HTS. We tested the effects of compounds in a commercial library on the MDM2/p53 interaction with above established assay. The library contains 2148 compounds, including five known inhibitors of MDM2/p53, Nutlin-3, Nutlin-3a, Nutlin-3b, YH239-EE and MI773. The activity of each compound was determined by assessing the enrichment of mMDM2 in the p53-containing droplets. Most compounds did not inhibit the recruitment of mMDM2. However, all five known MDM2 inhibitors (Fig. 3a, the 5 red dots) significantly reduced the mCherry signal and were the most effective inhibitors among all the compounds tested. Treatment with the five known inhibitors resulted in mCherry signals well over four standard deviations below the mean, which indicates that the screening process is effective. In addition, we found another compound, VER (abbreviated for VER155008, Fig. 3a, the green dot), which strongly attenuated mMDM2 enrichment, suggesting that it is a potential MDM2/p53 inhibitor. Fluorescence images confirmed that these inhibitors decreased the enrichment of mMDM2 in the phase-separated droplets (Fig. 3b).

**Figure 3.**
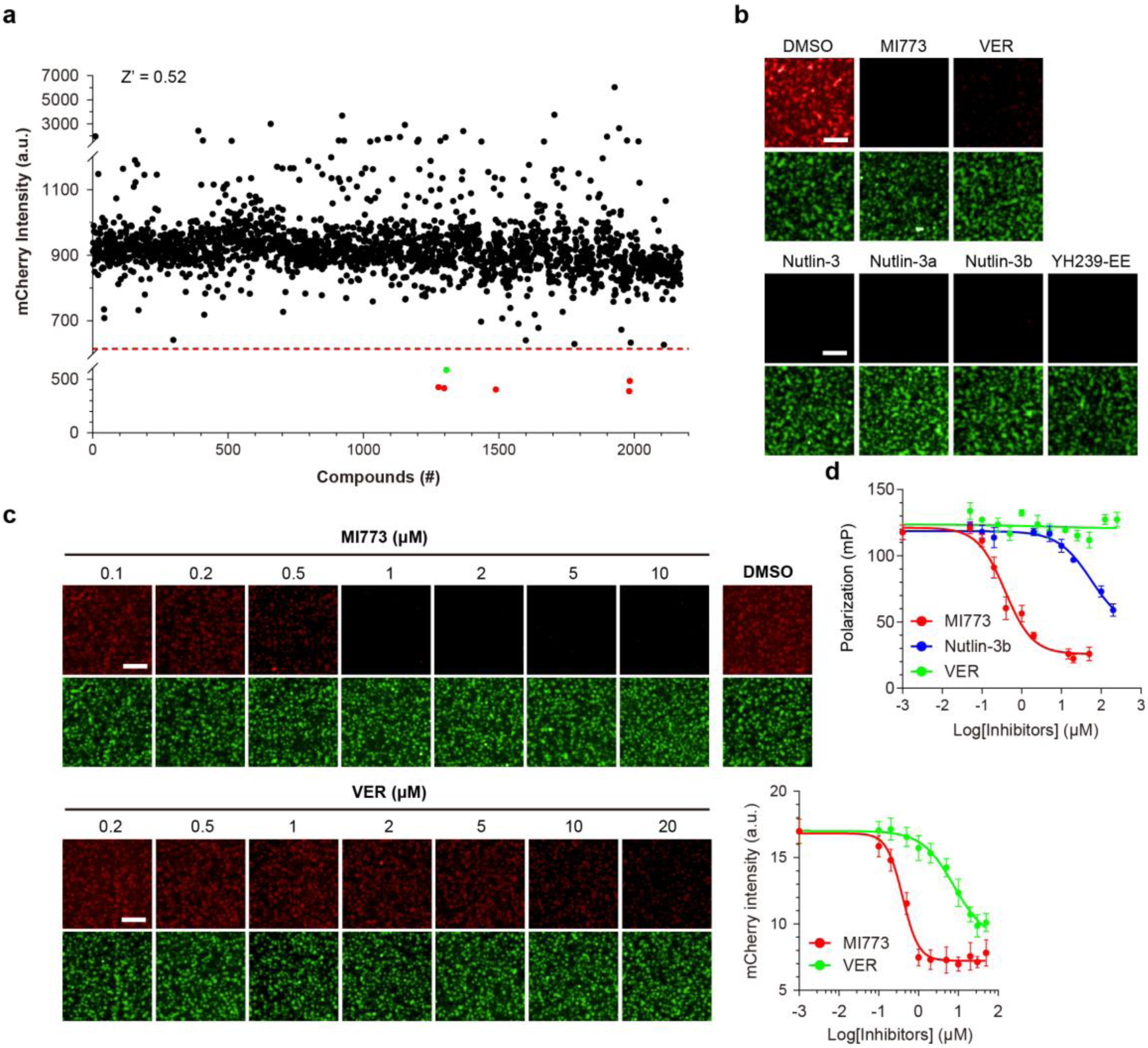
A high-throughput screen targeting the p53/MDM2 interaction. (a) An HTS targeting p53/MDM2 was performed using a commercial compound library, Selleck-2148. The concentration of each compound was 20 µM. The potential effect of each compound was evaluated by quantifying mCherry signal in the phase-separated green droplets. For visualization purposes, a red dashed line is arbitrarily drawn at MEAN - 4 × standard deviations. The 6 dots below the red dashed line are the 5 known MDM2 inhibitors (red dots) and an unknown MDM2 inhibitor, VER (green dot). All compounds in the library were numbered sequentially. YH239-EE (#1277), MI773 (#1298), VER (#1306), Nutlin-3 (#1489), Nutlin-3a (#1981), Nutlin-3b (#1983). (b) Fluorescence images showing the inhibition of the 6 screening hits on mMDM2 recruitment. (c) Dose-response analysis of MI773 and VER to confirm their inhibitory effects on mMDM2 recruitment. Representative fluorescence images for these analyses are shown. Dose-response curves (red for MI773 and green for VER) are plotted. (d) Competitive fluorescence polarization assay to evaluate the inhibitory effect of VER (green), MI773 (red), and Nutlin-3b (blue) on the mMDM2/p53 interaction. All scale bars, 10 μm.

The Z factor is a critical quality control parameter for HTS assays and it was 0.52 for our screen (Fig. 3a), which suggests that our assay is excellent for HTS^32^. An improved Z value (Z’ = 0.72) was obtained when we evaluated the enrichment of MDM2-SNAP (Fig. S2d) labeled with SNAP-Surface 546 in the droplets with a microplate reader after removal of the surrounding bulk solution (Fig. S4).

Detailed dose-response analysis confirmed the inhibitory activities of these screening hits (Fig. 3c and S5). Next, a competitive fluorescence polarization (FP) assay using a high affinity fluorescent probe P4 ^33^ was performed to confirm the inhibitory effect of the newly identified compound VER on the MDM2/p53 interaction. Two known MDM2 inhibitors, MI773 and Nutlin-3b, were used as controls. The control inhibitors caused an apparent decrease in polarization as their concentration increased, but VER failed to do so (Fig. 3d). This result suggested that CEBIT is more sensitive than FP. Collectively, these data revealed that CEBIT has the potential for drug discovery via HTS.

### Tailoring CEBIT to probe reader-based molecular modifications

Regulated chemical modifications (including methylation, acetylation, phosphorylation) of various biomacromolecules, such as proteins, RNAs, and DNAs, are extremely important for all aspects of biology. Aberrant modifications, and the associated enzymes are attractive therapeutic targets. However, efficient HTS assays that are generally applicable to the discovery of drugs modulating these enzymes are still lacking. These modifications can often be recognized by specific effector proteins via their reader domains. We postulated that CEBIT can be adopted for examining these modifications and the activities of the associated modifying enzymes. Our strategy is to confine specific reader domains in phase-separated compartments via fusion to the scaffold and use the modification-dependent recruitment of cognate substrates as the readout for the enzymatic activities of interest.

Here we used methylation of biomacromolecules (proteins, RNAs, and DNAs) as examples to test our idea. Protein lysine methylation plays important roles in various biological pathways. For example, methylation of the histone H3 lysine 9 (H3K9me) is a hallmark of heterochromatin in cells. We wonder whether this modification can be detected with CEBIT. To this end, we fused the human HP1β chromo domain (CD), a well-known reader domain of H3K9me^34–36^, to (gSH3)_14_, resulting in a composite scaffold (gSH3-CD)_14_ (Fig. 4a and S2e). We also fused H3K9 peptides (carrying different levels of lysine methylation) with the PDZ ligand, KKETPV, to create H3K9-KKETPV (abbreviated to H3K9-PV). Our strategy was to create droplets via LLPS of (gSH3-CD)_14_ in the presence of (gPRM)_14_ and then to determine the enrichment of H3K9-PV within the droplets by the further recruitment of mCherry labeled PDZ (Fig. 4a). We first used mono methylated H3K9-PV peptide (H3K9(me1)-PV) to test our design. As indicated, (gSH3-CD)_14_ and (gPRM)_14_ formed green droplets; however, a very weak mCherry signal was observed in the presence of mPDZ (Fig. 4b). To enhance the signal, mPDZ was multimerized by fusing with the hexameric protein BsHfq, resulting in (mPDZ)_6_ (Fig. S1 and S2e). Compared with mPDZ, (mPDZ)_6_ significantly enhanced the mCherry signal within the green droplets (Fig. 4b), which was confirmed by quantification (Fig. 4c). Hence, (mPDZ)_6_ was used for subsequent H3K9 methylation related studies.

**Figure 4.**
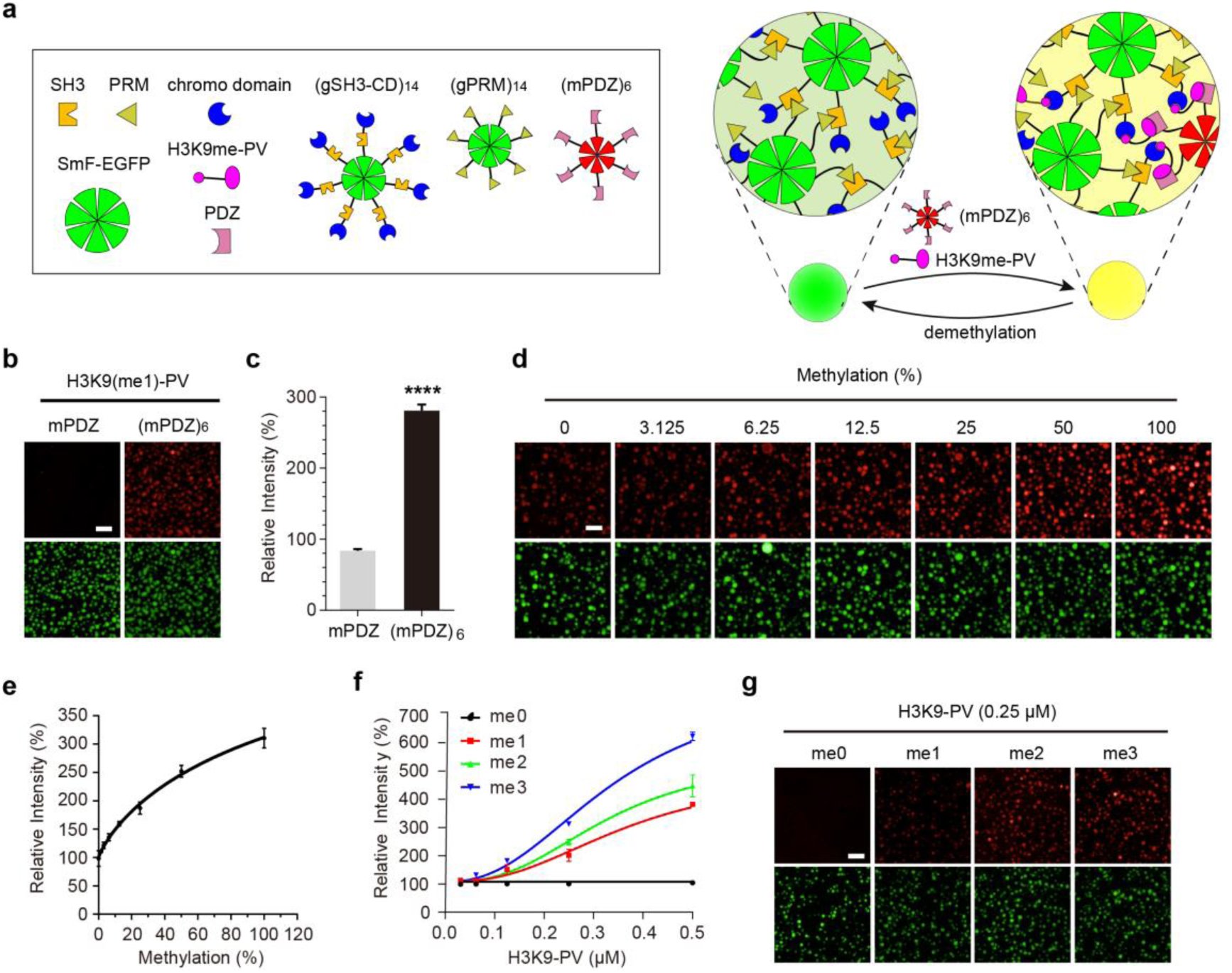
Application of CEBIT for probing molecular methylation. (a) Schematic diagram of CEBIT for detecting methylation of histone 3 at lysine 9. The individual modules are shown in the left box. The HP1β chromo domain (CD) is fused with (gSH3)_14_, resulting in the composite scaffold (gSH3-CD)_14_ which undergoes LLPS with (gPRM)_14_. As shown on the right, the methylation status of a synthetic H3K9 peptide can be assessed by its enrichment in the CD-containing compartments. For visualization, the H3K9 peptide is fused with the PDZ ligand KKETPV (H3K9-PV) which further binds to (mPDZ)_6_. (b-c) Synthetic mono-methylated H3K9-PV peptide (H3K9(me1)-PV; 1 μM) was used in the methylation assay system containing 1 μM (gPRM)_14_, 1 μM (gSH3-CD)_14_ and 1 μM mPDZ or (mPDZ)_6_. Then enrichment of H3K9(me1)-PV in phase-separated green droplets was analyzed. Fluorescence images (b) and quantification of H3K9(me1)-PV peptide enrichment in the phase-separated droplets (c) are shown. Normalized mCherry signal were shown. **** p<0.0001. (d-e) Mixtures of H3K9(me0)-PV and H3K9(me1)-PV at various ratios (total concentration, 1 μM) were used in the assay system containing 1 μM (gPRM)_14_, 1 μM (gSH3-CD)_14_, and 1 μM (mPDZ)_6_. For each mixture, enrichment of mCherry signal in the phase-separated green droplets were assessed by both imaging (d) and quantification (e). Normalized mCherry signal were shown. (f-g) H3K9-PV peptides with different levels of methylation were used in the methylation assay system which contained 1 μM (gPRM)_14_, 1 μM (gSH3-CD)_14_, and 1 μM (mPDZ)_6_. Enrichment of these peptides in the phase-separated green droplets was assessed. (f) Quantification of mCherry signal in the phase-separated droplets. Normalized mCherry signal were shown. (g) Fluorescence images for the 4 different peptides at 0.25 μM are presented. **** p<0.0001. All scale bars, 10 μm.

Next, we used mixtures of H3K9(me0)-PV and H3K9(me1)-PV with increasing ratios of the latter to mimic the methylation reaction process. As the proportion of H3K9(me1)-PV increased, the mCherry signal increased in parallel (Fig. 4d-e). We next analyzed the effect of the methylation status of the H3K9-PV peptide (from me0 to me3) andfound thathigher levels of methylation resultedin strongermCherry signals(Fig. 4f-g), which was consistent with the binding preference of HP1β chromo domain^37, 38^.

We subsequently studied the methylation of DNA and RNA with CEBIT. The recognition of methylated CpG by the MBD domain of human MBD1 (which binds m^5^C on DNA) and of methylated RNA by the YTH domain of human YTHDC1 (which binds N-m^6^A on RNA) were taken as examples. In this experiment, we used another set of phase separation scaffolds of linear topology, polySUMO/polySIM^19^. The clients were ROX-labeled DNA or RNA oligonucleotides carrying two MBD- or YTH-recognition motifs, of which 0, 1 or 2 were methylated (designated DNA(0me)/RNA(0me), DNA(1me)/RNA(1me), and DNA(2me)/RNA(2me)). Their corresponding recognition domains, MBD and YTH, were fused to GFP-labeled tandem pentameric SUMO3, g(SUMO3)_5_ as the composite scaffolds, which assembled liquid compartment with a hexameric SUMO3 interacting motif, SIM_6_ (Fig. S6a) via LLPS. We reasoned that the methylation status of DNA and RNA could be assessed by their partition in the MBD- or YTH-enriched compartments, respectively. g(SUMO3)_5_-MBD and SIM_6_ formed green droplets via multivalent SUMO3/SIM interaction. Both DNA(1me) and DNA(2me) were appreciably enriched within these droplets, and the latter was more strongly enriched than the former (Fig. S6b). This observation was further confirmed by quantification of ROX intensity in the droplets (Fig. S6b). Additionally, when mixtures of DNA(0me) and DNA(2me) with various ratios but at a constant total concentration were used, the enrichment of ROX signal in MBD-containing droplets was proportional to the ratio of DNA(2me) (Fig. S6c). For RNA, identical experiments were performed and similar results were achieved (Fig. S6d-e). Collectively, these results showed that CEBIT can effectively probe the methylation of biomacromolecules.

### CEBIT-based HTS for SUV39H1 inhibitors

SUV39H1 is a critical methyltransferase that catalyzes the transfer of methyl groups to H3K9^39^. We took SUV39H1 as an example to assess the capacity of CEBIT to screen for drugs that modulate various enzymes. The activity of SUV39H1 was evaluated using the assay system shown in Figure 4a. The substrate was H3K9(me0)-PV, and SUV39H1 activity was assessed by enrichment of mCherry signal in the droplets. Indeed, the mCherry signal was enhanced as the concentration of SUV39H1 increased in the *in vitro* methylation reaction (Fig. 5a). Prolonging the reaction time also increased the mCherry signal (Fig. 5b). In contrast, methylation of H3K9(me0)-PV by SUV39H1 was largely abolished by a potent SUV39H1 inhibitor, chaetocin, and a feedback inhibitor for generic methyltransferase reactions, S-adenosine homocysteine (SAH). The IC50 values were 1.6 and 56 μM for chaetocin and SAH, respectively (Fig. 5c-d).

**Figure 5.**
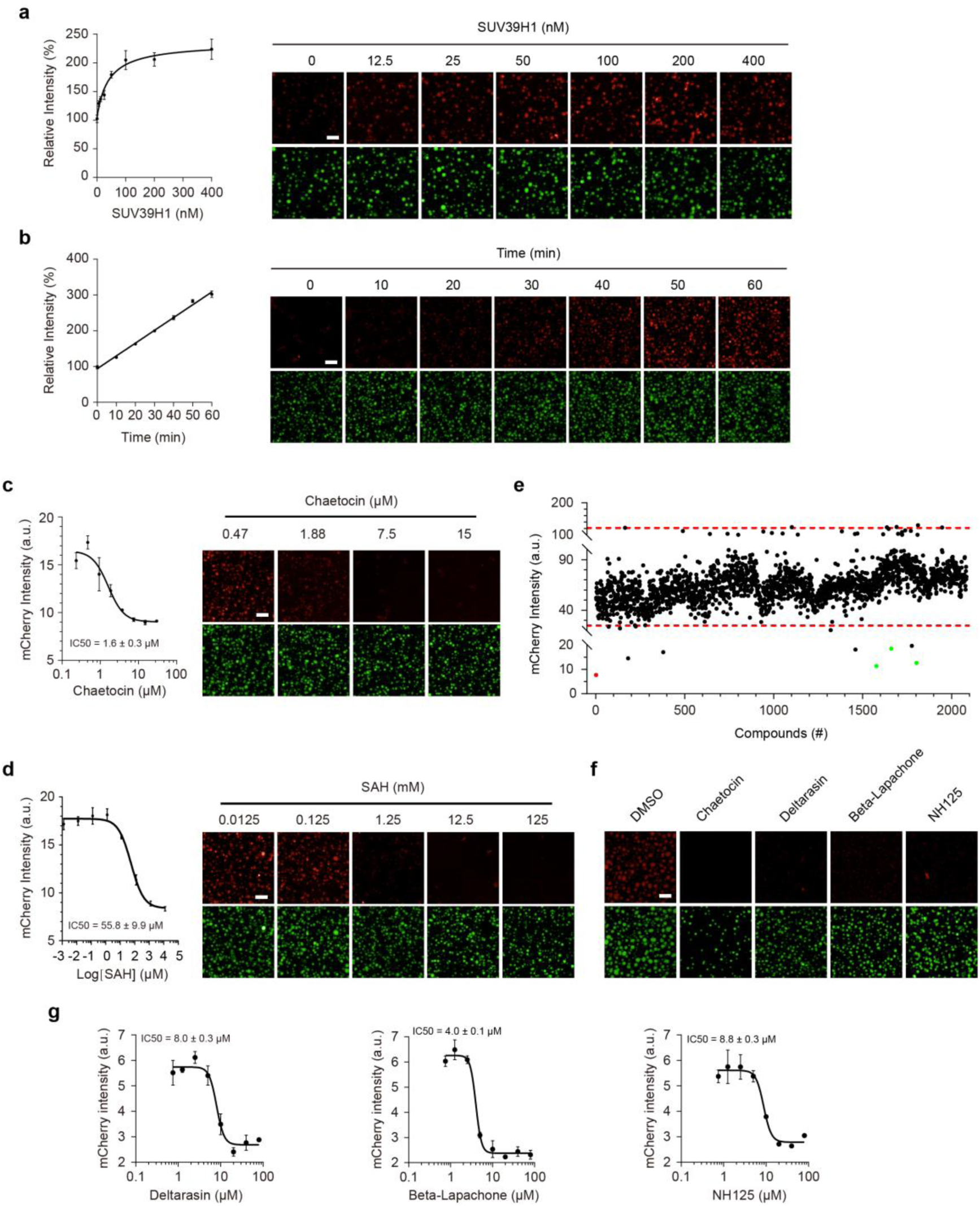
Discovery of SUV39H1 inhibitors via CEBIT based high-throughput screening. (a) *In vitro* methylation of H3K9(me0)-PV peptide (8 μM) by various amounts of SUV39H1 was conducted and the reaction products were subjected to the methylation assay system shown in Fig. 4a. Normalized mCherry signal in the phase-separated green droplets was plotted against SUV39H1 concentration (left panel). Representative fluorescence images are shown (right panel). (b) *In vitro* methylation of H3K9(me0) PV (8 μM) by SUV39H1 (0.4 μM) was conducted and then the reaction products were detected by imaging. The time-course of SUV39H1 activity was evaluated by plotting normalized mCherry signal against reaction time. Fluorescence images are presented. (c-d) Analysis of the *in vitro* methylation of H3K9(me0)-PV (8 μM) by SUV39H1 (0.4 μM) in the presence of various doses of the SUV39H1 inhibitors chaetocin (c) and SAH (d). The reaction products were assessed with the indicated assay, and mCherry intensity was plotted against inhibitor concentration. (e-f) High-throughput screening for discovery of SUV39H1 inhibitors. (e) For each compound in the library, enrichment of SUV39H1-methylated peptide substrate in the phase-separated green droplets was assessed according to the intensity of the mCherry signal. Screening hits are indicated by colored dots. The red dot represents chaetocin (#1) and the three green dots represent Deltarasin (#1578), Beta-Lapachone (#1661) and NH125 (#1803). (f) Fluorescence images showing the inhibition of enrichment by these compounds. (g) Dose-response analysis of screening hits to confirm their inhibitory effect on SUV39H1 activity. mCherry signal in phase-separated compartments was evaluated. All scale bars, 10 μm.

We next explored its potential for the discovery of SUV39H1 inhibitors via HTS. We screened the Selleck-2148 compound library in which chaetocin was added as a positive control. Treatment with chaetocin resulted in the lowest mCherry signal in the phase separated compartments (Fig. 5e, red dot, and 5f). A few compounds from the library also caused low mCherry signals (Fig. 5e-f). We selected three of them, Deltarasin, Beta-Lapachone, and NH125 (Fig. 5e, green) for further testing. Their inhibition of SUV39H1 activity was assessed by detailed dose-response analysis *in vitro* and their IC50 values were within the single-digit µM range (Fig. 5g). These results demonstrated that CEBIT worked well for *in vitro* HTS-based discovery ofdrugs that modulateSUV39H1 activity, suggesting a general application of CEBIT to other methyltransferase enzymes as well as enzymes for other molecular modifications, such as phosphorylation, acetylation, ubiquitinylation.

## Discussion

In this study, we have developed a HTS based drug discovery method called CEBIT which stands for <u>c</u>ompartmentalization of <u>e</u>nhanced <u>b</u>iomolecular interactions in test tubes. It’s generally amenable to determining diverse modification-dependent and -independent BIs and associated enzymes *in vitro,* and hence discovering therapeutic drugs to modulate these interactions and enzymatic activities via HTS. Our study showed that CEBIT is highly versatile and easy to use, it will become a powerful tool for drugs development.

CEBIT is distinct from existing *in vitro* PPI detection methods in two respects: 1) it is based on, and hence is analogous to, the general architecture of the prevalent membraneless organelles in cells; and 2) the actual readout is based on the recruitment of nanometer-sized molecules into micron-sized compartments.

Many membraneless organelles assembled via LLPS^13–16^ and the complex compositions of these assemblies are generally divided into two groups: scaffold and client^19^. This architecture revealed two distinct strategies to study various PPIs. In principle, the PPI of interest can serve as either the scaffold or the client. In the former setup, both partners of the PPI are multimerized via fusion to homo-oligomeric proteins and the appearance of phase-separated puncta serves as a readout for the interaction^20, 21, 40^. This strategy is adopted by two recent work, Fluoppi^20^ and SPARK^21^. In the latter setup, the recruitment of a client into its binding partner enriched compartments serves as a readout for the interaction, as revealed by our method, CEBIT. The former strategy has some major limitations. Phase separation of multimerized PPI is controlled by several factors, such as valence, concentration, stoichiometry and the binding affinity between PPI of interest. If any of the factors is below the threshold for LLPS, no puncta can be observed. Therefore, it’s less efficient and less reliable to use the former strategy for assaying various PPIs. In contrast, it’s free for CEBIT to select robust model scaffolds to drive the formation of phase separated compartments, such as SmF fused model PPIs (SH3/PRM, PDZ/PDZ ligand, SUMO3/SIM) and polySUMO/polySIM, these multivalent model interactions provide much more stable platforms to study BIs of interest. Additionally and importantly, multimerization cannot be easily achieved for many PPI components due to the insolubility of target proteins by fusion to homo-oligomeric protein. However, we can choose to implement the components of PPI of interest in monovalent fashion by using linear multivalent scaffolds (such as SH3_4_-PRM_4_^17^ and polySUMO-polySIM^19^) with CEBIT. For at least these reasons, we believe CEBIT is more general, easier, and more reliable than Fluoppi to study various PPIs, just by connecting one partner of the PPI to the scaffold and the other labeled with fluorophore as client. Fluoppi was claimed to have the potential for drug screening, however, no HTS campaign has been performed to characterize its behavior. Our study has provided solid experimental evidence to demonstrate that CEBIT is very amenable for drug discovery via HTS. SPARK, a phase separation based kinase reporter, is designed to visualize the dynamics of kinase activity in vivo by using similar strategy to Fluoppi. The kinase substrate and consensus binding protein are multimerized by respective fusion with homo-oligomeric protein HO-tag3 and HO-tag6. Upon kinase activation, phase separated droplets formed by multivalent interaction between phosphorylated substrate and its binding protein. Actually, similar problems for SPARK exist as Fluoppi. For example, the phosphorylation level of the kinase substrate is very low beneath its threshold for LLPS at the initial stage of kinase activation, there will be no cellular puncta. Our study has provided an easy and general way to investigate diverse biomacromolecular modifications and associated enzymes by taking biomolecular methylation (including protein, DNA and RNA) and SUV39H1 as examples.

SPR, FP and FRET are three of the most extensively used HTS technologies for drug discovery in vitro. SPR is sensitive and versatile for detection of diverse biomolecular interactions, however, many factors can interfere with the outcomes of SPR. Our work suggested that CEBIT is very robust for studying diverse interactions and tolerates the factors that interfere SPR. As the readout of CEBIT is the enrichment of fluorophore labeled client in its binding partner containing compartment, its signal dose not affected by the difference in molecular mass between interaction pairs. Compared to FRET, CEBIT exhibited good signal window for study diverse PPIs (PDZ/PDZ ligand, p53/MDM2, Smac/XIAP, H3K9me /chromo domain), what’s more, the signal can further be enhanced by increasing the valence of the client (Fig. 4b, S6b and S6d). Additionally, CEBIT based assay is simple and efficient, from HTS assay development, to detection and data analysis.

CEBIT is highly versatile and has great scope for adaptation and improvement as far as its applicability and efficiency are concerned. It is conceivable that by fusing Streptavidin to the phase separation scaffolds, CEBIT can be used to detect any BIs with one component of the interaction pair biotinylated (unpublished data). Besides *in vitro* studies, this method can also be applied to study diverse PPIs *in vivo* (see the accompanying manuscript for details) and to identify regulators of these PPIs via HTS in cells. What’s more, CEBIT is likely to analyze the interactions between membrane receptors and their ligands by creating membrane-attached receptor-enriched compartments. Other important applications will likely be developed.

## Material and methods

### Reagents

All modified DNAs and RNAs were provided by Zixi Biotech Company (Beijing, China). Peptides used in this study were chemically synthesized by GL Biochem Ltd (Shanghai, China). The fluorescent probe SNAP-Surface Alexa Fluor 546 (#S9132S) was from NEB Biotech. The chemical library Selleck-2148 for HTS and compounds for the dose-response assay were provided by the drug facility of Tsinghua University. Other small molecules were purchased from Selleck.

### Plasmid construction

All the plasmids used in this study were constructed following normal molecular cloning techniques and fragments of interest was confirmed by sequencing. All the SmF-derived composite scaffold proteins were created on the basis of SmF-EGFP or SmF-mCherry, which was generated by sequentially inserting SmF and EGFP or mCherry into pRSF-Duet1 (Invitrogen) with an N-terminal His6 tag, except (Smac-SH3)_14_ and (SH3-p53)_14_. To produce multivalence, interacting module domains (including SH3/PRM, PDZ/PDZ ligand and SUMO3/SIM) were respectively fused to SmF-EGFP/mCherry to create multimeric composite scaffold proteins such as SmF-EGFP-SH3 (abbreviated to (gSH3)_14_) and SmF-mCherry-PRM (abbreviated to (mPRM)_14_).

To generate (Smac-SH3)_14_ and (SH3-p53)_14_, two more backbone constructs were first generated in pRSF-Duet1: pHMS1 was created by sequentially inserting His6, MBP, SUMO tag (cleaved by ULP1 protease) and SmF; and pHMS2 was created by inserting His6, MBP, and SUMO-Smac. To construct (SH3-p53)_14_, a PCR fragment of SH3-p53 was ligated to pHMS1. To generate (Smac-SH3)_14_, a PCR fragment of SmF-SH3 was inserted into pHMS2.

In this study, (gSH3)_14_ and (gPRM)_14_ were the most extensively used scaffolds. To study protein-protein interactions of interest, one member of the interaction pair was fused to (gSH3)_14_ as the composite scaffold, and the other was fused with an N-terminal mCherry tag as the client. To multimerize mCherry-PDZ, a hexamer protein BsHfq was fused at the N terminus.

Another set of scaffold proteins used in this study, polySUMO and polySIM, were modified from the generous gifts from Professor MK Rosen (UTSW, USA). PCR fragments of (SUMO3)_5_ and SIM_6_ were inserted into pRSF-Duet1 fused with an N-terminal His6 tag. GFP was further fused to the C-terminus of (SUMO3)_5_, resulting in g(SUMO3)_5_. To detect nucleic acid methylation, the indicated DNA or RNA binding domains (MBD for methylated DNA and YTH for methylated RNA) were fused to g(SUMO3)_5_ to generate the composite scaffold proteins.

A full-length cDNA encoding the methyltransferase SUV39H1 was amplified by PCR and fused with an N-terminal His6-MBP tag in pET-Duet1 (Invitrogen).

### Protein expression and purification

Generally, recombinant proteins were expressed in *Escherichia coli* BL21 (DE3) cells in LB medium with 0.5 mM IPTG at 18℃ for 12-24 hours unless noted. The bacteria were lysed by sonication in buffer (50 mM Tris pH 8.0, 500 mM NaCl, 10% glycerol and 1 mM PMSF), then proteins were purified using nickel-NTA agarose beads (GE Healthcare, UK) followed by ion exchange chromatography and final by size exclusion chromatography with KMEI buffer (150 mM KCl, 1 mM EGTA, 1 mM MgCl_2_, 10 mM imidazole pH 7.4, 1 mM TCEP and 10% glycerol). Some proteins were not amenable to ion exchange, and therefore this purification step was omitted.

For (SH3-p53)_14_ and (Smac-SH3)_14_, the purification was performed by sequentially using nickel-NTA agarose beads, anion exchange, cleavage by ULP1, a tandem MBP-NTA column to remove the N-terminal tags, and finally size exclusion chromatography with buffer containing 150 mM KCl, 1 mM EGTA, 1 mM MgCl_2_, 10 mM imidazole pH 7.4, 1 mM TCEP and 10% glycerol. All proteins were rapidly frozen with liquid nitrogen and stored at -80℃.

PolySUMO-derived proteins [including g(SUMO3)_5_-YTH/MBD] and SIM_6_ were individually expressed and affinity-purified with nickel-NTA beads followed by size exclusion chromatography in buffer containing 150 mM KCl, 1 mM MgCl_2_, 1 mM EGTA, 20 mM HEPES pH 7.0, 1 mM DTT (the pH is critical for the phase separation of polySUMO and polySIM as revealed by the work of Banani et al^19^).

To achieve active enzyme, SUV39H1 and HP1β were co-expressed in *Escherichia coli* BL21 (DE3) cells and induced with 0.5 mM IPTG at 18℃ for 16 hours. The complex of SUV39H1 and HP1β was affinity-purified with nickel-NTA beads followed by size exclusion chromatography as described above.

### Size exclusion chromatography coupled with multiangle light scattering (SEC-MALS)

The analysis was performed as previously described^16^. The protein samples were centrifuged at 12,000g for 10 min to remove any protein precipitation and then loaded into a gel filtration column (Superdex 200 Increase 10/300GL, GE Healthcare) coupled with an 18 angle light scattering detector (DAWN HELEOS II, Wyatt) and an Optilab DSP interferometric refractometer (Wyatt) in buffer composed of 150 mM KCl, 1 mM EGTA, 1 mM MgCl_2_, 10 mM imidazole pH 7.4. Data were analyzed by ASRTA (Wyatt).

### Client recruitment and inhibition assays

SmF-derived scaffold proteins, client proteins and inhibitors were mixed together at desired concentrations in buffer KMEI (150 mM KCl, 1 mM EGTA, 1 mM MgCl_2_, 10 mM imidazole pH7.4) in a total volume of 20 µL and incubated at 4℃ for more than 2 hours to allow the phase-separated droplets gradually sinking down to the bottom of the wells in a 384-well microplate. Images were then taken by high-content microscopy (Opera Phenix, Perkin Elmer, USA) and the fluorescence intensity in the phase-separated droplets was analyzed using the microscopy-linked software Harmony4.8. Images in Figures 1 and 2b were taken and analyzed by confocal microscopy (Nikon A1RSi, Japan).

### PolySUMO- and polySIM-based partitioning assay

PolySUMO- and polySIM-based partitioning assays were conducted in buffer containing 20 mM HEPES pH 7.0, 150 mM KCl, 1 mM EGTA, 1 mM MgCl_2_, 1 mM DTT. The phase-separated scaffolds and recruited clients were mixed thoroughly followed by incubation at 4℃ for more than 2 hours. Images were taken and analyzed by high-content microscopy.

### Partitioning assay of SNAP tagged proteins

A mixture of 1 μM (Smac-SH3)_14_, 1 μM (gPRM)_14_, 2 μM XIAP-SNAP, and 1 μM SNAP-Surface 546 fluorophore was incubated with XIAP inhibitor GDC-0152 at the indicated concentration in KMEI buffer supplemented with 2% DMSO at 4℃ overnight for efficient labeling of XIAP-SNAP. Enrichment of XIAP-SNAP in phase-separated green droplets was assessed by imaging and evaluation of the SNAP-Surface 546 signal.

Similarly, 1 μM (SH3-p53)_14_, 1 μM (gPRM)_14_, 2 μM MDM2-SNAP and 1 μM SNAP-Surface 546 fluorophore were incubated with 5 μM MI773 in KMEI buffer supplemented with 2% DMSO at 4℃ overnight. As expected, (SH3-p53)_14_ and (gPRM)_14_ formed green droplets via LLPS, into which MDM2-SNAP was recruited. Recruitment of MDM2-SNAP was inhibited by MI773. The phase-separated green droplets were isolated by cautious removal of the surrounding bulk solution after centrifugation. Enrichment of fluorescently labeled MDM2-SNAP in the droplets was assessed by microplate reader via measurement of SNAP-Surface 546 intensity. The Z factor was calculated with fluorescent intensity of DMSO (0% inhibition) and 5uM MI773 (100% inhibition).

### Competitive fluorescence polarization (FP) assay

The FP assay was performed as described by Czarna et al^33^. The probe was a p53-derived synthetic peptide P4 modified with FITC. The interacting protein was recombinant human MDM2 (residues 10-110) with an N-terminal His6 tag. The FP assay was performed with 5 nM FTIC-P4 and 80 nM His-MDM2 in the presence of different concentrations of MI773, Nutlin-3b and VER. All assay components were mixed thoroughly in the buffer (40 mM Tris pH 8.0, 150 mM NaCl, 1 mM EDTA and 5% DMSO) with a total volume of 10 μL. The FP value was measured by using a versatile microplate reader (EVISION, Perkin Elmer, USA) after 30 min incubation at room temperature.

### *In vitro* methylation and detection of H3K9 peptide

The synthetic peptide substrate H3K9(me0)-PV (8 μM) was *in vitro*-methylated by SUV39H1-HP1β complex in the presence of 20 μM SAM. The reaction was conducted in buffer containing 50 mM Tris pH 8.4, 5 mM MgCl_2_, 4 mM DTT by incubation at room temperature for 2 hours and ended by heating at 95℃ for 10 min to inactivate the enzyme. Subsequently, 5 μL of the reaction product was analyzed in the CEBIT system consisting of 1 μM (gSH3-CD)_14_, 1 μM (gPRM)_14_, 1 μM (mPDZ)_6_ in a final volume of 20 μL in KMEI. After incubation for 2 hours at 4℃, images were taken by high-content microscopy. mCherry signal in the phase-separated droplets was analyzed. To determine the time-course of enzyme activity, the reaction was stopped by quick-freezing with liquid nitrogen followed by heating at 95℃ for 10 min to inactivate the enzyme. Then the methylation of H3K9(me0)-PV peptide was assessed as indicated.

### High-throughput screening

The assay system for screening compounds that disrupt the p53/MDM2 interaction was comprised of 0.5 μM (gSH3-p53)_14_, 0.5 μM (gPRM)_14_ and 1 μM mMDM2 in KMEI buffer supplemented with 2% DMSO in a total volume of 20 μL in a 384-well microplate (Greiner Bio-One, Cat. 781090). High-throughput screening was performed using compounds in a commercial library (Selleck-2148). This library contains 2148 small molecules, which was supposed to contain 6 reported MDM2 inhibitors (Nutlin-3, Nutlin-3a, Nutlin-3b, YH239-EE, MI773, and RG7388). However, the solution corresponding to RG7388 in the stock library failed to compete p53/MDM2 after repeated trials while separately purchased RG7388 potently inhibited p53/MDM2 using CEBIT. We concluded that RG7388 in the library used was misplaced and 5 known MDM2 inhibitors (Nutlin-3, Nutlin-3a, Nutlin-3b, YH239-EE, and MI773) actually existed in the library. Each compound was used at a concentration of 20 μM. After thoroughly mixing all the components (scaffolds, client proteins and compounds), the microplates were incubated at 4℃ for overnight before imaging and data collection.

In the screen for inhibitors of the methyltransferase SUV39H1, compounds in the Selleck-2148 library were used at a final concentration of 30 μM. Each compound was added to an *in vitro* methylation reaction comprising 8 μM H3K9(me0)-PV, 0.4 μM SUV39H1 and 20 μM SAM. The mixture was incubated at room temperature for 2 hours before the reaction was stopped by heating at 60℃ for 30min. Subsequently, 5 μL product from the reaction was assessed using the indicated methylation assay system. After incubation at 4℃ for 2 hours, images were taken by high content microscopy and data were collected.

**Table 1.**
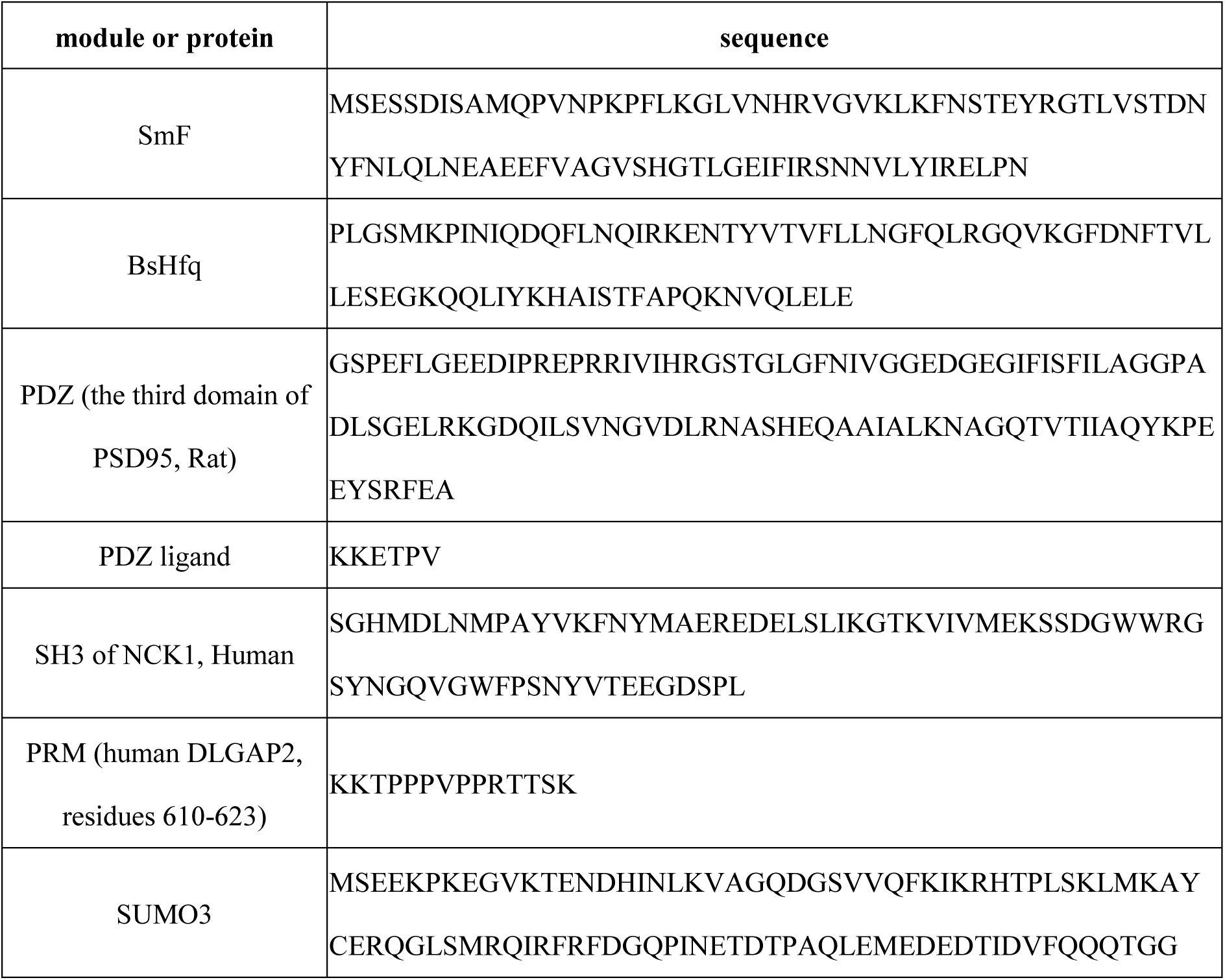
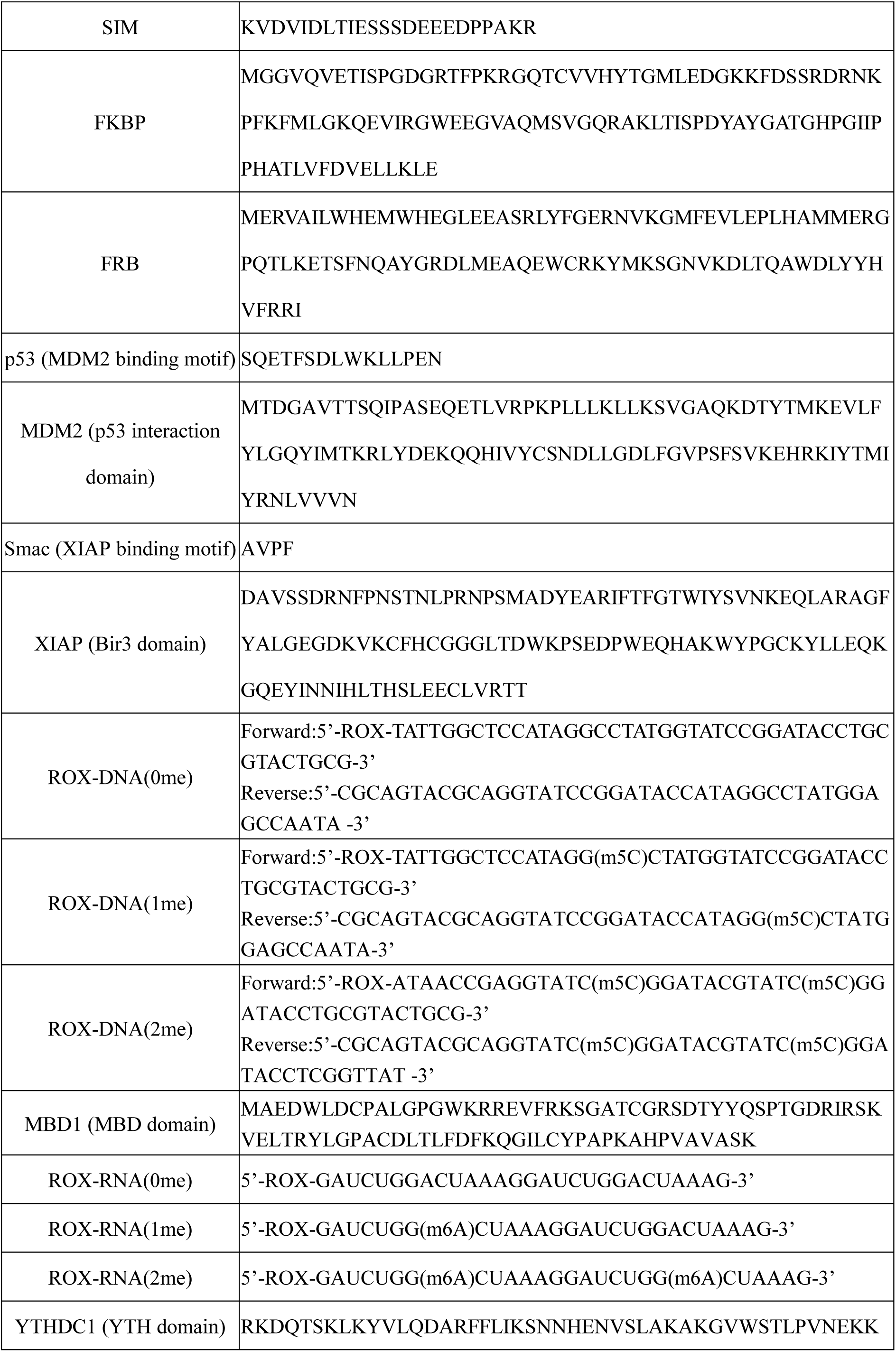
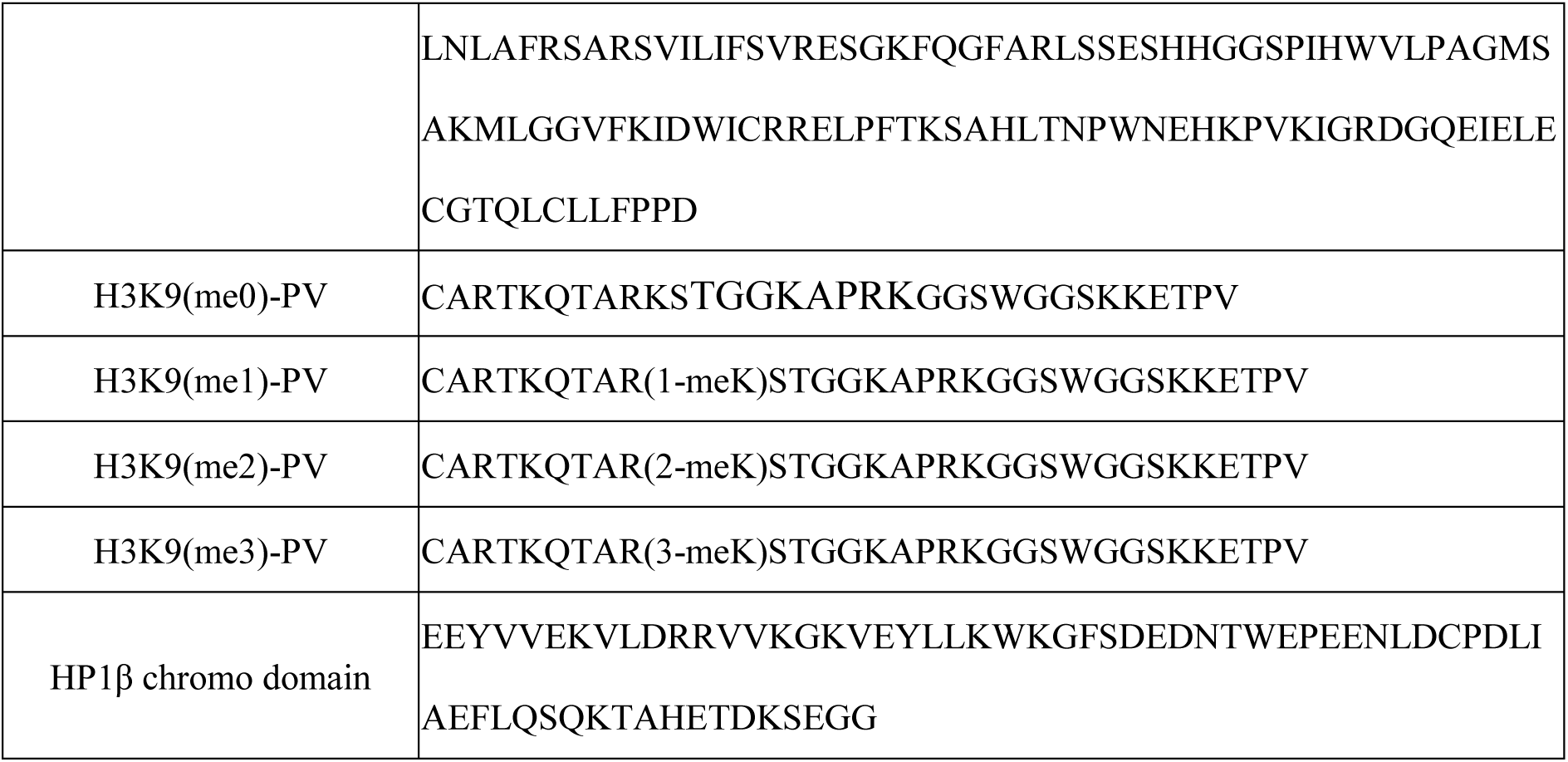
Modules and proteins used in this study

## Acknowledgements

This work was supported by grants from the National Key R&D Program (2019YFA0508403 to P.L.) and the Natural Science Foundation of China (31871443 to P.L.; 31800637 to J.W.).

## Author contributions

P.L. conceived and supervised the project; M.Z., W.L., J.L., R.W., L.W., S.F., W.S., and J.H. carried out the experiments; M.Z. and P.L. wrote the manuscript with help from W.L. and J.L..

## Supplementary Figures

**Figure S1.**
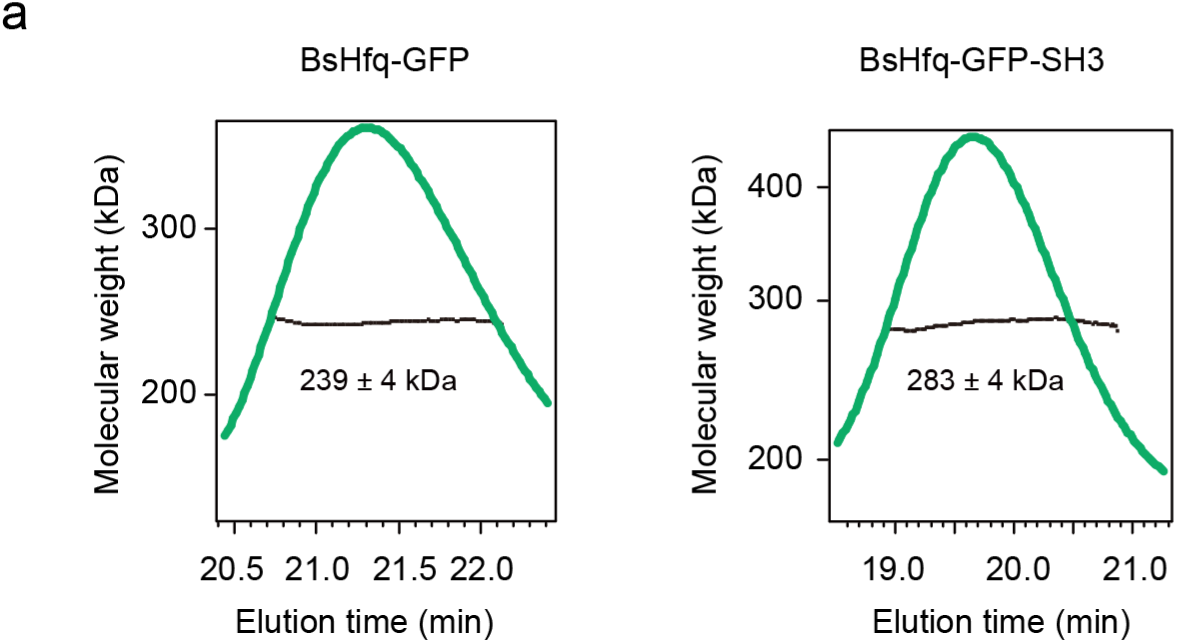
SEC-MALS analysis of BsHfq-fused multivalent proteins. SEC-MALS analysis of BsHfq-GFP (theoretical molecular mass 238 kDa) and BsHfq-GFP-SH3 (theoretical molecular mass 293 kDa), showing that both fusion proteins remained hexameric.

**Figure S2.**
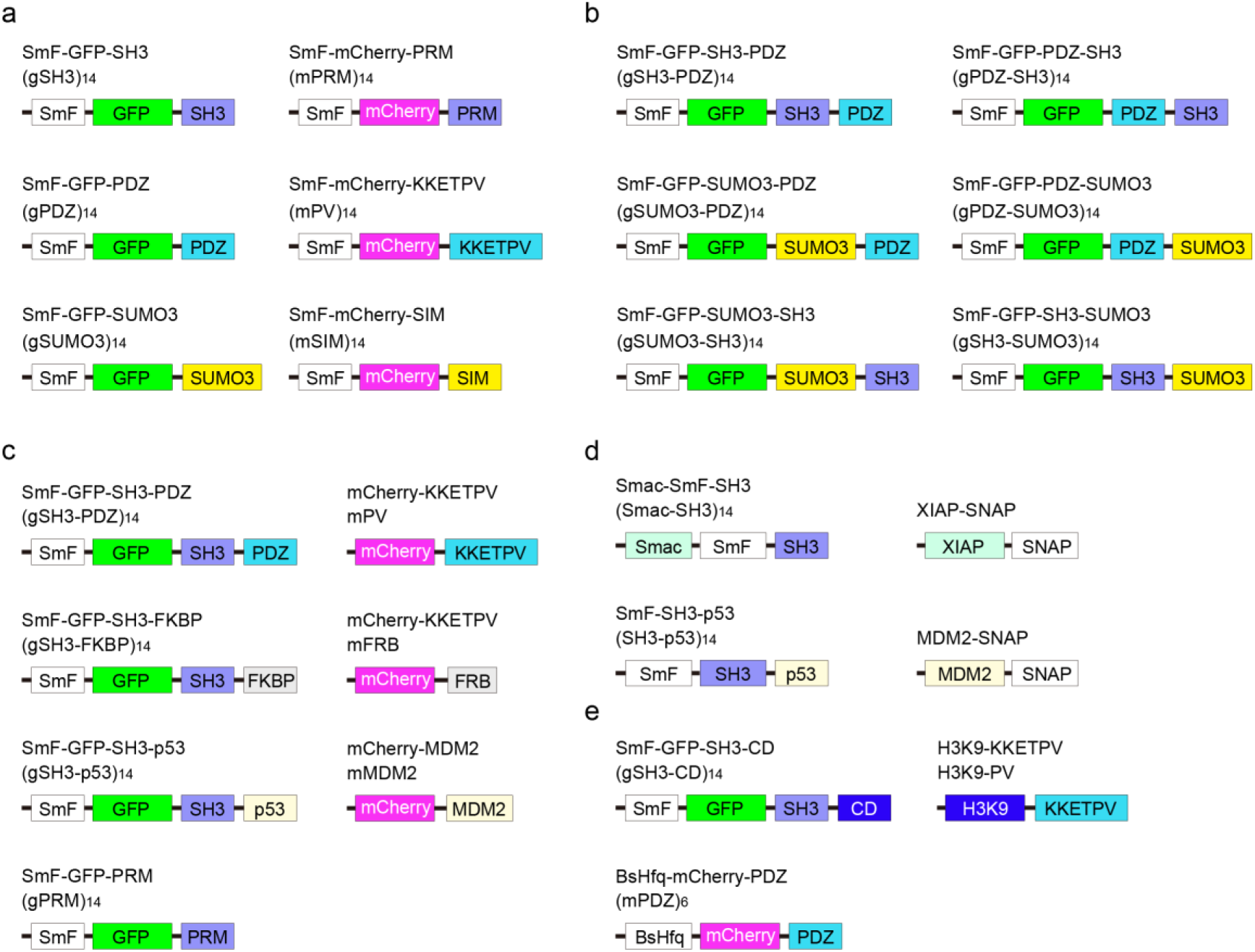
Domain organization of SmF-fused multivalent scaffolds and clients used in this study. (a-e) Domain architecture of the scaffold proteins and clients used in Fig. 1b (a), Fig. 1d (b), Fig. 2 (c), Fig. S3 and S4 (d) and Supplementary Fig. 4 (e).

**Figure S3.**
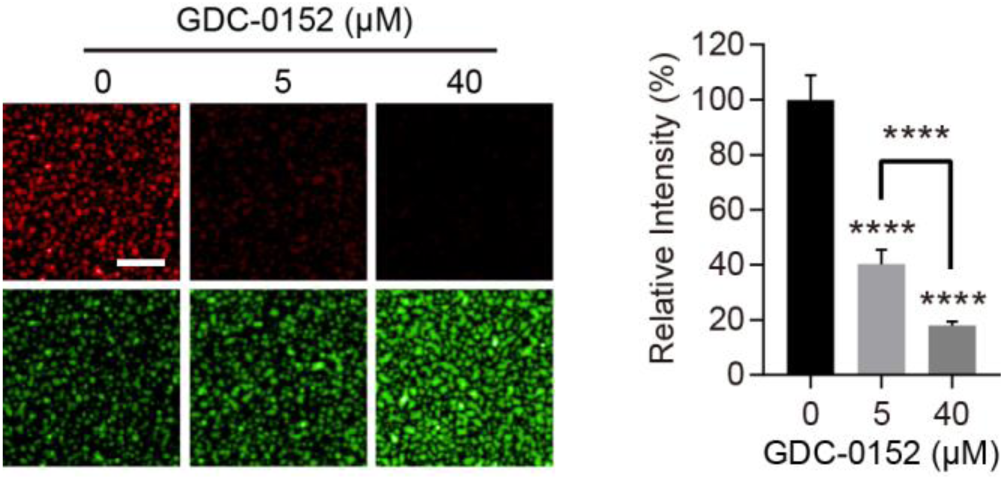
Recruitment of XIAP-SNAP into Smac-enriched phase-separated compartments. (a-b) Enrichment of SNAP-Surface 546-labeled XIAP-SNAP (2 μM) into phase-separated green droplets formed by (Smac-SH3)_14_ and (gPRM)_14_ (1 μM each). The enrichment was challenged by the XIAP inhibitor GDC-0152. Fluorescence images and quantification of SNAP-Surface 546 in the green droplets were shown.

**Figure S4.**
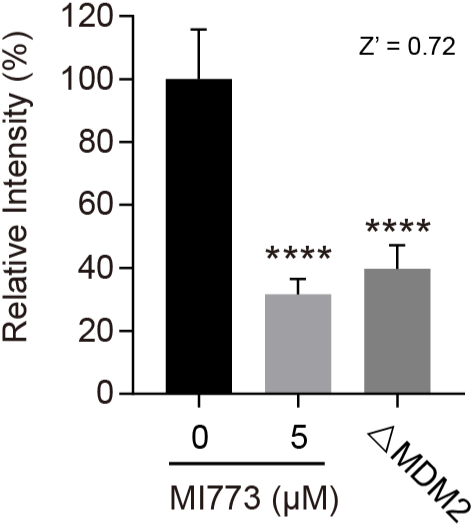
An improved Z factor of CEBIT based MDM2/p53 interaction assay. SNAP-Surface 546-labeled MDM2-SNAP was recruited into p53-enriched droplets formed by (SH3-p53)_14_ and (gPRM)_14_. This recruitment was completely inhibited by 5 μM MI773. MDM2-SNAP^SNAP-Surface 546^ enrichment in phase-separated droplets was evaluated by microplate reader after removal of the surrounding bulk solution. The samples without addition of MDM2-SNAP (△MDM2) was used as positive control. Z factor was calculated with fluorescent intensities of samples treated with DMSO (0% inhibition) and 5 μM MI773 (100% inhibition).

**Figure S5.**
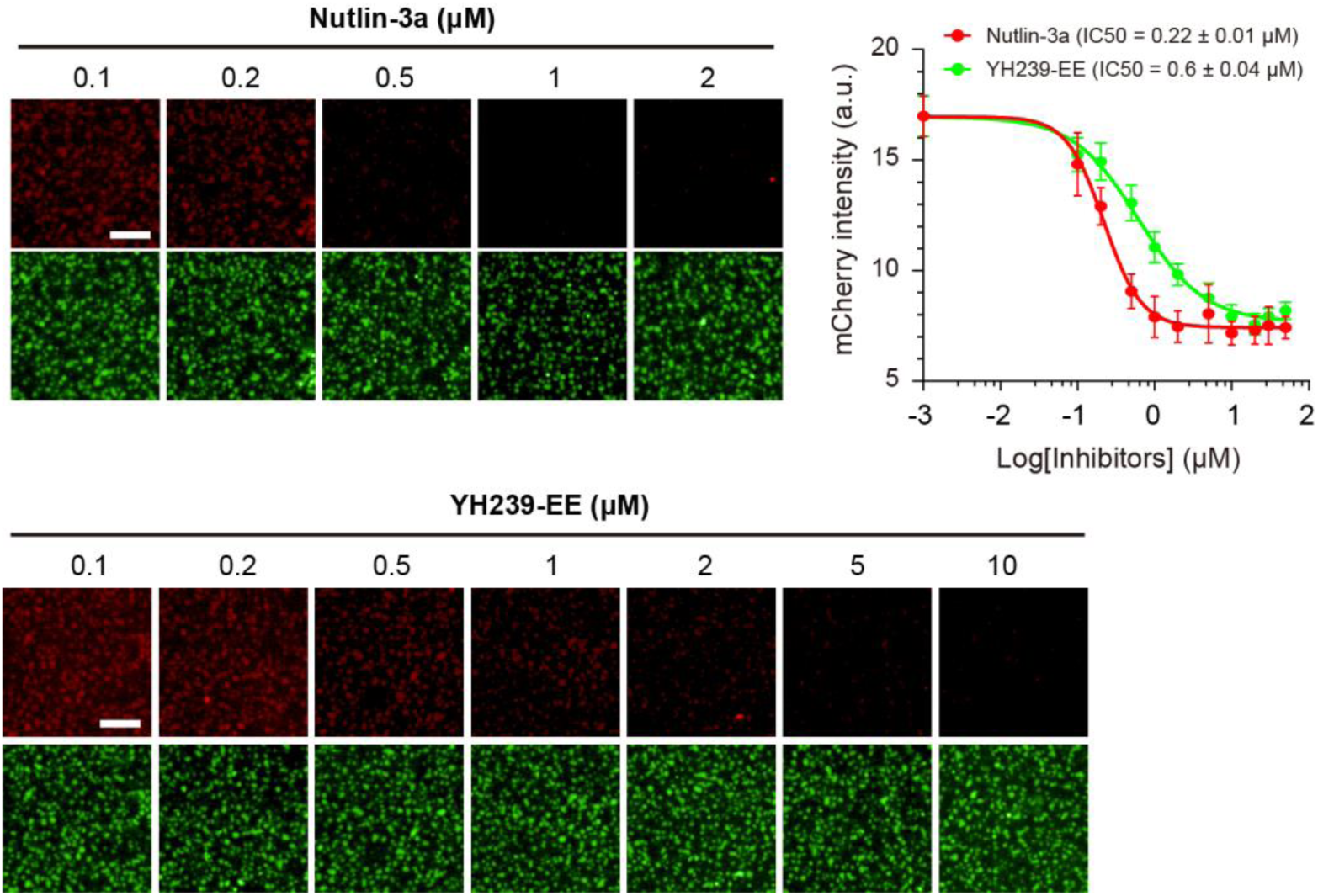
Dose-response analysis of the HTS hits for inhibition of mMDM2 recruitment. Dose-response analysis of Nutlin-3a and YH239-EE for inhibition of MDM2/p53 interaction with the assay system containing 0.5 μM (gSH3-p53)_14_, 0.5 μM (gPRM)_14_ and 1 μM mMDM2. Representative fluorescence images and dose-response curves were shown.

**Figure S6.**
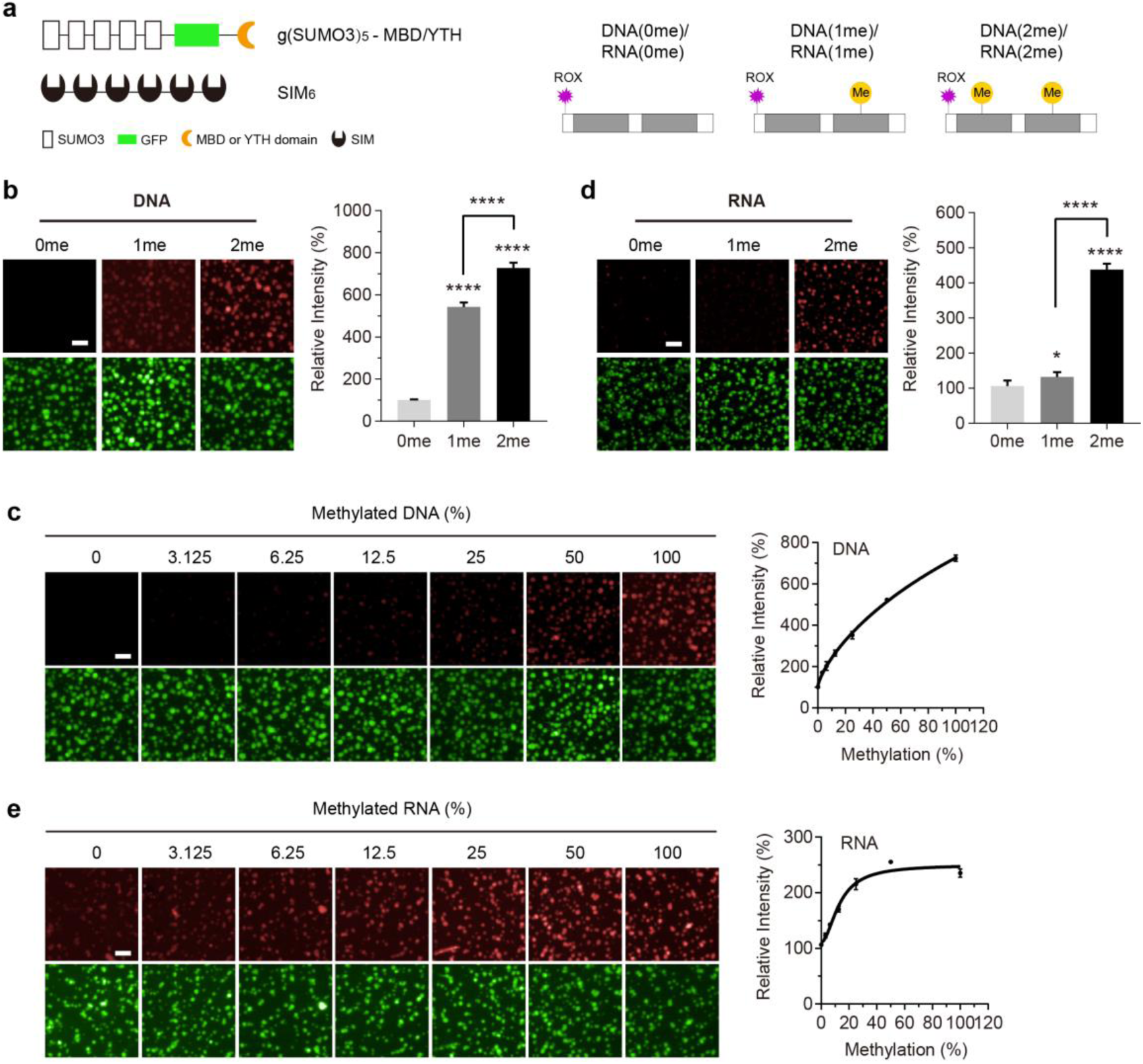
Assessment of nucleic acid methylation with CEBIT. (a) Schematic of the assay system for DNA and RNA methylation. The scaffold proteins for inducing LLPS are SIM_6_ and g(SUMO3)_5_-MBD (for DNA methylation) or g(SUMO3)_5_-YTH (for RNA methylation). g(SUMO3)_5_ refers to (SUMO3)_5_-EGFP. The recruited clients are fluorophore-labeled methylated DNA or RNA oligos. The DNA or RNA oligos consist of two repeated protein-binding modules and carry 0, 1 or 2 methyl groups. (b) Partitioning of ROX-labeled DNA (0.2 μM) carrying 0, 1 or 2 methyl groups into compartments induced by g(SUMO3)_5_-MBD and SIM_6_ (2 μM each). Fluorescence images and quantification of DNA enrichment in phase-separated green droplets were presented. Normalized ROX signal were shown. (c) Mixtures of ROX-labeled DNA(0me) and DNA(2me) (total concentration, 1 μM) at different ratios were subjected to the DNA methylation assay system comprising 2 μM g(SUMO3)_5_-MBD and SIM_6_. Methylation-dependent DNA enrichment into the phase separated droplets was assessed. Representative images and normalized ROX signal in the droplets are shown. (d) Recruitment of ROX-labeled RNA (0.2 μM) carrying 0, 1 or 2 methyl groups into phase-separated droplets formed by g(SUMO3)_5_-YTH and SIM_6_ (2 μM each). Fluorescence images and normalized ROX signal in phase-separated droplets are shown. (e) Mixtures of ROX-labeled RNA(0me) and RNA(2me) (total concentration, 1 μM) were assessed by their partitioning into phase-separated compartments containing the YTH domain (2 μM each of scaffold proteins). Representative images and normalized ROX signal in the green droplets are shown.

## References

1. Ryan, D.P. & Matthews, J.M. Protein-protein interactions in human disease. Current Opinion in Structural Biology 15, 441–446 (2005).

2. Kuzmanov, U. & Emili, A. Protein-protein interaction networks: probing disease mechanisms using model systems. Genome Medicine 5, 37 (2013).

3. Ivanov, A.A., Khuri, F.R. & Fu, H. Targeting protein-protein interactions as an anticancer strategy. Trends Pharmacol Sci 34, 393–400 (2013).

4. Vassilev, L.T. et al. In vivo activation of the p53 pathway by small-molecule antagonists of MDM2. Science 303, 844–848 (2004).

5. Flygare, J.A. et al. Discovery of a potent small-molecule antagonist of inhibitor of apoptosis (IAP) proteins and clinical candidate for the treatment of cancer (GDC-0152). J Med Chem 55, 4101–4113 (2012).

6. Macarron, R. et al. Impact of high-throughput screening in biomedical research. Nature Reviews Drug Discovery 10, 188–195 (2011).

7. Giannetti, A.M. From experimental design to validated hits a comprehensive walk-through of fragment lead identification using surface plasmon resonance. Methods Enzymol 493, 169–218 (2011).

8. Renaud, J.P. et al. Biophysics in drug discovery: impact, challenges and opportunities. Nature Reviews Drug Discovery 15, 679–698 (2016).

9. Zhou, M., Li, Q. & Wang, R. Current Experimental Methods for Characterizing Protein-Protein Interactions. ChemMedChem 11, 738–756 (2016).

10. Sipova-Jungova, H. et al. Biomolecular charges influence the response of surface plasmon resonance biosensors through electronic and ionic mechanisms. Biosens Bioelectron 126, 365–372 (2019).

11. Syafrizayanti, Betzen, C., Hoheisel, J.D. & Kastelic, D. Methods for analyzing and quantifying protein-protein interaction. Expert Review of Proteomics 11, 107 (2014).

12. Janzen, William & nbsp Screening Technologies for Small Molecule Discovery: The State of the Art. Chemistry & Biology 21, 1162–1170 (2014).

13. Brangwynne, C.P. et al. Germline P granules are liquid droplets that localize by controlled dissolution/condensation. Science 324, 1729–1732 (2009).

14. Feric, M. et al. Coexisting liquid phases underlie nucleolar sub-compartments. Cell 165, 1686–1697 (2016).

15. Amandine, M. et al. Phase Separation by Low Complexity Domains Promotes Stress Granule Assembly and Drives Pathological Fibrillization. Cell 163, 123–133 (2015).

16. Zeng, M. et al. Phase Transition in Postsynaptic Densities Underlies Formation of Synaptic Complexes and Synaptic Plasticity. Cell 166, 1163–1175.e1112 (2016).

17. Li, P. et al. Phase transitions in the assembly of multivalent signalling proteins. Nature 483, 336–340 (2012).

18. Banani, S.F., Lee, H.O., Hyman, A.A. & Rosen, M.K. Biomolecular condensates: organizers of cellular biochemistry. Nature Reviews Molecular Cell Biology 18, 285–298 (2017).

19. Banani, S.F. et al. Compositional Control of Phase-Separated Cellular Bodies. Cell 166, 651–663 (2016).

20. Watanabe, T. et al. Genetic visualization of protein interactions harnessing liquid phase transitions. Scientific Reports 7, 46380 (2017).

21. Qiang, Z. et al. Visualizing Dynamics of Cell Signaling InVivo with a Phase Separation-Based Kinase Reporter. Molecular Cell 69, 347 (2018).

22. Sun, D., Wu, R., Zheng, J., Li, P. & Yu, L. Polyubiquitin chain-induced p62 phase separation drives autophagic cargo segregation. Cell Research 28, 405–415 (2018).

23. Collins, B.M. et al. Homomeric ring assemblies of eukaryotic Sm proteins have affinity for both RNA and DNA. Crystal structure of an oligomeric complex of yeast SmF. J Biol Chem 278, 17291–17298 (2003).

24. Tatsuhiko, S. et al. Crystal structure of Hfq from Bacillus subtilis in complex with SELEX-derived RNA aptamer: insight into RNA-binding properties of bacterial Hfq. Nucleic Acids Research 40, 1856–1867 (2012).

25. Saro, D. et al. A thermodynamic ligand binding study of the third PDZ domain (PDZ3) from the mammalian neuronal protein PSD-95. Biochemistry 46, 6340–6352 (2007).

26. Banaszynski, L.A., Liu, C.W. & Wandless, T.J. Characterization of the FKBP.rapamycin.FRB ternary complex. Journal of the American Chemical Society 127, 4715–4721 (2005).

27. Kussie, P.H. et al. Structure of the MDM2 oncoprotein bound to the p53 tumor suppressor transactivation domain. Science 274, 948–953 (1996).

28. Wang, S., Zhao, Y., Aguilar, A., Bernard, D. & Yang, C.Y. Targeting the MDM2-p53 Protein-Protein Interaction for New Cancer Therapy: Progress and Challenges. Cold Spring Harb Perspect Med 7, a026245 (2017).

29. Liu, Z. et al. Structural basis for binding of Smac/DIABLO to the XIAP BIR3 domain. Nature 408, 1004 (2000).

30. Wang, S. et al. SAR405838: an optimized inhibitor of MDM2-p53 interaction that induces complete and durable tumor regression. Cancer Res 74, 5855–5865 (2014).

31. Ding, Q., et al. Discovery of RG7388, a potent and selective p53-MDM2 inhibitor in clinical development. J Med Chem 56, 5979–5983 (2013).

32. Zhang, J.H., Chung, T.D.Y. & Oldenburg, K.R. A Simple Statistical Parameter for Use in Evaluation and Validation of High Throughput Screening Assays. Journal of Biomolecular Screening 4, 67–73 (1999).

33. Anna, C. et al. High affinity interaction of the p53 peptide-analogue with human Mdm2 and Mdmx. Cell Cycle 8, 1176–1184 (2009).

34. Bannister, A.J. et al. Selective recognition of methylated lysine 9 on histone H3 by the HP1 chromo domain. Nature 410, 120–124 (2001).

35. Lachner, M., O’Carroll, D., Rea, S., Mechtler, K.,. & Jenuwein, T., Methylation of histone H3 lysine 9 creates a binding site for HP1 proteins. Nature 410, 116–120 (2001).

36. Jacobs, S.A. & Sepideh, K. Structure of HP1 chromodomain bound to a lysine 9-methylated histone H3 tail. Science 295, 2080–2083 (2002).

37. Fischle, W. et al. Molecular basis for the discrimination of repressive methyl-lysine marks in histone H3 by Polycomb and HP1 chromodomains. Genes Dev 17, 1870–1881 (2003).

38. Hughes, R.M., Wiggins, K.R., Sepideh, K. & Waters, M.L. Recognition of trimethyllysine by a chromodomain is not driven by the hydrophobic effect. Proceedings of the National Academy of Sciences of the United States of America 104, 11184–11188 (2007).

39. Rea, S. et al. Regulation of chromatin structure by site-specific histone H3 methyltransferases. Nature 406, 593–599 (2000).

40. Nakamura, H., DeRose, R. & Inoue, T. Harnessing biomolecular condensates in living cells. J Biochem 166, 13–27 (2019).

